# Evaluation of sample preservation and storage methods for metaproteomics analysis of intestinal microbiomes

**DOI:** 10.1101/2021.07.20.453169

**Authors:** Angie Mordant, Manuel Kleiner

## Abstract

A critical step in studies of the intestinal microbiome using meta-omics approaches is the preservation of samples before analysis. Preservation is essential for approaches that measure gene expression, such as metaproteomics, which is used to identify and quantify proteins in microbiomes. Intestinal microbiome samples are typically stored by flash freezing and storage at −80°C, but some experimental set-ups do not allow for immediate freezing of samples. In this study, we evaluated methods to preserve fecal microbiome samples for metaproteomics analyses when flash freezing is not possible. We collected fecal samples from C57BL/6 mice and stored them for 1 and 4 weeks using the following methods: flash-freezing in liquid nitrogen, immersion in RNAlater™, immersion in 95% ethanol, immersion in a RNAlater-like buffer, and combinations of these methods. After storage we extracted protein and prepared peptides for LC-MS/MS analysis to identify and quantify peptides and proteins. All samples produced highly similar metaproteomes, except for ethanol-preserved samples that were distinct from all other samples in terms of protein identifications and protein abundance profiles. Flash-freezing and RNAlater™ (or RNAlater-like treatments) produced metaproteomes that differed only slightly, with less than 0.7% of identified proteins differing in abundance. In contrast, ethanol preservation resulted in an average of 9.5% of the identified proteins differing in abundance between ethanol and the other treatments. Our results suggest that preservation at room temperature in RNAlater™, or an RNAlater-like solution, performs as well as freezing for the preservation of intestinal microbiome samples before metaproteomics analyses.

**Importance:** Metaproteomics is a powerful tool to study the intestinal microbiome. By identifying and quantifying a large number of microbial, dietary, and host proteins in microbiome samples, metaproteomics provides direct evidence of the activities and functions of microbial community members. A critical step for metaproteomic workflows is preserving samples before analysis because protein profiles are susceptible to fast change in response to changes in environmental conditions (air exposure, temperature changes, etc.). This study evaluated the effects of different preservation treatments on the metaproteomes of intestinal microbiome samples.

## Introduction

The intestinal microbiome is a highly diverse and metabolically active community that has profound effects on its host [1]. This complex community influences the health of its host by altering the availability of nutrients [2–5] and the host’s susceptibility to infection and disease [6, 7]. The intestinal microbiome is integral to proper host immune function [8–10] and host well-being [11, 12]. So far most studies have used DNA sequencing and taxonomy-based approaches to study the intestinal microbiome providing critical insights into taxonomic shifts in the community-related host genotype, diet, and disease state [13–16]. Taxonomic shifts, however, have been found to not always measure important functional shifts in the microbiome, as different taxa can perform the same function [17] and highly similar strains can perform different functions by encoding a few unique gene clusters [18]. Therefore the use of function-focused multi-omics approaches is essential for understanding the role of the intestinal microbiome in health and disease [19–22].

Metaproteomics is a valuable tool to study interactions in the intestinal microbiome and the microbiome’s influence on host health [23–25]. Metaproteomics allows for the identification and quantification of large numbers of microbial, dietary, and host proteins in microbiome samples in a high-throughput fashion [26, 26–28]. Because proteins are central to all biological processes, metaproteomics provides direct evidence of the activities and functions of microbial community members and their contributions to disease [29]. For example, metaproteomics revealed protein biomarkers of disease in Inflammatory Bowel Disease (IBD) and colorectal cancer and provided insights into the role of the microbiome in such diseases [30, 31]. In addition to quantifying differences in protein abundances between samples, metaproteomics can also be used to assess microbial community structure based on proteinaceous biomass [32–35] and track incorporation of specific substrates using stable isotope content of peptides [36–39].

Metaproteomics workflows are complex and variability can be introduced at every step, from sample preparation to data acquisition by LC-MS/MS and data processing [35]. There is no standardized workflow for metaproteomics of intestinal microbiome samples but some of the individual steps have been optimized in the past, such as protein extraction [40], database creation [41], and database searching [42]. A critical step that has not yet been optimized is the storage and preservation of samples before analysis. Adequate storage of samples is critical because exposure to environmental changes could induce changes in protein profiles of species in the samples and thus provide misleading study results. For example, exposing samples to air can strongly bias colorectal cancer studies because oxidative stress and enrichment of bacterial superoxide dismutase enzymes that will occur from air exposure are also characteristics of colorectal cancer in the intestinal tract [30]. Therefore, appropriately storing samples immediately upon collection helps to avoid post-collection changes in protein abundances. A suitable storage method should preserve the information contained in the microbiome at the time of sampling without introducing substantial bias.

Typically, microbiome samples are frozen immediately upon collection, with flash-freezing in liquid nitrogen followed by storage at −80° C. However, flash-freezing is not always possible, and very little is known about suitable alternatives to flash-freezing for the preservation of microbiome samples before metaproteomics analyses. For example, clinical or diet studies involving human subjects usually require the subjects to perform the sampling themselves at their homes where they do not have access to liquid nitrogen [43, 44]. It can be difficult to maintain sample integrity also in resource-limited fieldwork conditions [45] or out in a wild animal’s environment where there is no access to liquid nitrogen and cold storage [46]. Even in the laboratory, immediate freezing in liquid nitrogen can be difficult. One specific case is work with gnotobiotic animals, which are invaluable models to study and manipulate the microbiota in a controlled system. Gnotobiotic animals reside in isolators where everything (food, bedding, etc) entering the isolators is sterilized through autoclaving, irradiation, or strong chemicals before being introduced through a two-ended port [47]. Removing samples from the isolators causes long delays between sampling and sample storage, thus exposing samples to environmental changes (air exposure, temperature change, nutrient depletion, etc) before they can be adequately stored.

Several studies have been conducted to test the effects of preservation methods on nucleic acids, but the effects of such methods on proteins are poorly understood. Freezing at −80°C is thought to maintain sample integrity similar to fresh samples [48, 49]. The effectiveness of storage at room temperature is typically evaluated based on comparisons to frozen treatments. RNA*later*™ is a popular storage solution that has been shown to be effective at preserving DNA in gut microbiome studies, with negligible differences compared to freezing [50, 51]. Ethanol (95% or absolute) may also be suitable for the preservation of nucleic acids in microbiome samples before taxonomic profiling as long as it is used consistently [45, 46, 52]. However, there are studies in which these storage solutions significantly biased the downstream results [53], particularly in RNA sequencing studies [54] and thus great care should be taken when selecting a preservation method. While the effects of sample preservation on nucleic acids have been extensively studied, to the best of our knowledge, only two studies have investigated the effects of sample preservation on protein profiles. First, Saito and colleagues demonstrated that RNA*later*™ has the potential to preserve proteomes as effectively as immediate freezing [55]. Their results are promising, however, their study was performed on a single marine microorganism (cyanobacterium *Synechococcus* WH8102) and thus does not indicate whether RNA*later*™ would preserve samples as complex as those from the intestinal microbiome. Furthermore, their study was performed in earlier stages of proteomics when replication was expensive and effort-intensive. For that reason, the researchers only included technical replication and therefore they could not assess the robustness of RNA*later*™ in terms of within-treatment consistency. Second, Hickl and colleagues observed vast differences in the identifications and relative abundances of proteins from human fecal samples depending on the preservation and storage procedure applied to the samples [56]. They tested two preservation and storage methods: a flash-freezing-based approach “FF” and RNAlater, “RL”. The first method “FF”, consisted of flash-freezing in liquid nitrogen followed by storage at −80°C, cryomilling, and storage at −20°C for 16 h while immersed in RNA*later*ICE®. The “RL” method simply consisted of immersion in refrigerated RNA*later*™ for 6 h. They found less than 50% overlap in protein identifications between the two treatments. Of the overlapping proteins, they found roughly 2,000 proteins to significantly differ in abundance by more than 1.5 fold between the two treatments. The majority of the differences they observed were attributed to taxonomy. For example, class Actinobacteria represented about 20% of the composition of the FF samples while Actinobacteria only made up less than 2% of the composition of the RL samples. However, one cannot attribute the differences Hickl et al. observed to a specific aspect of the preservation and storage process because of the many variables in the sample processing. For example, the flash-frozen samples were cryomilled which could favor lysis of gram-positive bacteria, such as Actinobacteria, and may explain the large difference in the relative abundance of this taxon [57].

This study aimed to (1) compare the effects of different sample preservation methods on intestinal microbiome metaproteomes, (2) evaluate comparable aspects of sample processing by limiting the number of variables, (3) assess within-treatment variability, and (4) evaluate the methods over a longer period of preservation/storage time.

## Materials and Methods

### Preparation of fecal master mix and preservation treatments

Fresh fecal pellets were collected from 12 conventional 5-month old C57BL/6 mice obtained from the Jackson Laboratory. To remove inter-individual variation as a variable, the pellets were pooled and homogenized using a spatula to make a fecal master mix. The master mix was split into aliquots of 8 mg each. The aliquots were either resuspended in 200 μL of a preservation solution and stored at room temperature (~22° C) in the dark, or were flash-frozen using liquid nitrogen and stored at −80° C. The preservation treatments are discussed in detail below. In brief, we tested 6 preservation treatments: flash-freezing in liquid nitrogen followed by storage at −80° C (“FF” treatment); preservation at room temperature in RNA*later*™ (RNA*later*™ Stabilization Solution, Invitrogen; “R” treatment); a combination of RNA*later*™ and flash freezing (“RF” treatment); an RNA*later*™-like preservation buffer (“NAP buffer”) as described in [58] (“N” treatment); this same “NAP buffer” autoclaved (“AN” treatment); and 95% ethanol (“E” treatment). We tested the effectiveness of the treatments at preserving microbiome samples over two storage durations: 1 week and 4 weeks. We prepared four replicate samples per treatment and time point, with each replicate being an 8 mg aliquot of the fecal master mix described above.

#### Flash-freezing and storage at −80° C (“FF”)

Immediate freezing followed by storage at −80° C is the method most frequently used to preserve biological specimens and is regarded as the “gold-standard” approach. Fouhy and colleagues observed that immediate freezing retains information similar to fresh samples in a 16S rRNA gene amplicon sequencing experiment of healthy human fecal samples [48]. The only significant differences they observed between frozen and fresh samples were in the relative abundances of the genera *Faecalibacterium* and *Leuconostoc;* however, the differences were subtle and may be attributable to a batch effect in DNA extraction rather than sample preservation. The effectiveness of storage solutions used at room temperature is typically evaluated based on comparisons to frozen treatments [55].

#### Immersion in RNAlater™ and storage at room temperature (“R”)

RNA*later*™ Stabilization Solution is a popular storage reagent. Its effectiveness can be attributed to its ability to quickly permeate tissue to stabilize and protect RNA. RNAl*ater*™ is effective at preserving nucleic acids for intestinal microbiome studies, with negligible bias compared to freezing [50]. RNA*later*™ has the potential to preserve proteins because its main component is ammonium sulfate, and ammonium sulfate precipitates proteins which can later be re-solubilized without degradation. Saito et al. demonstrated that RNAl*ater*™ is effective at preserving the proteome of the marine cyanobacterium *Synechococcus* WH8102 [55]. In our study, we immersed samples in RNA*later*™ (Invitrogen) in a 1:10 sample: solution ratio and then stored them at room temperature (~22° C) in the dark.

#### Immersion in RNAlater™ and flash-freezing followed by storage at room temperature (“RF”)

To determine whether the use of a storage solution makes a difference as compared to storing samples dry at −80° C, we immersed RF samples in RNA*later*™ (Invitrogen) and flash-froze the tubes in liquid nitrogen before storing them at −80° C. Observed differences between R and RF samples would provide evidence regarding the effects of freezing on sample integrity.

#### Immersion in NAP buffer and storage at room temperature (“N”)

The major limitation of RNA*later*™ is its high cost. It has been demonstrated that RNA*later*™-like buffers work as effectively as the commercially available solution [59]. Menke and colleagues even argue that their RNAlater-like solution called “Nucleic Acid Preservation (NAP) buffer” outperformed commercial RNA*later*™ in preserving DNA for 16S rRNA gene amplicon sequencing experiments based on comparisons with immediately-frozen controls [58]. “NAP buffer” was included as a treatment in this study and was prepared as previously described by Camacho-Sanchez et al. [59]. Briefly, 1.5 L of NAP buffer (pH 5.2) contained 935 ml of ultrapure water, 700 g of ammonium sulfate, 25 ml of 1 M sodium citrate, and 40 ml of 0.5 M ethylenediaminetetraacetic acid (EDTA). We prepared the solution fresh two days before the experiment. We immersed the samples in the NAP buffer solution in a 1:10 sample: solution ratio before storing them at room temperature (~22° C) in the dark.

#### Immersion in Autoclaved NAP buffer and storage at room temperature (“AN”)

RNA*later*™ and RNAlater-like buffers do not need to be autoclaved because their chemical composition inhibits the growth of contaminants. Manufacturers of RNA*later*™ recommend against autoclaving the reagent. However, in some cases, such as when working with gnotobiotic isolators, the solution needs to be autoclaved to prevent the introduction of microorganisms into the isolators. We tested an autoclaved version of the NAP buffer as an additional treatment to simulate real experimental conditions with gnotobiotic isolators. The same solution described above as the N treatment, from the same batch, was autoclaved (60 min at 121.5° C) two days before the start of the experiment. We immersed the samples in the autoclaved NAP buffer solution in a 1:10 sample: solution ratio before storing them at room temperature (~22° C) in the dark.

#### Immersion in 95% Ethanol and storage at room temperature (“E”)

Alcohol preservation is a common method in which biological specimens are preserved by dehydration. Hale and colleagues [46] found that absolute ethanol worked as well as immediate freezing of DNA for preserving samples prior to metagenomics analysis. The effectiveness of ethanol as a preservation treatment depends on its concentration. Sinha et al. [60] observed low stability of microbial DNA when preserved in 70% ethanol. Saito et al. [55] observed that 90% was not ideal for the preservation of the proteome of the marine cyanobacterium *Synechococcus* WH8102 as only ~75% of the proteins were recovered as compared to flash-freezing. Because the organism studied by Saito et al. [55] is very different from the intestinal microbiome, we included ethanol (95%) as a treatment. We prepared 95% ethanol by mixing pure anhydrous (200 proof/100%) ethyl alcohol (Koptec) with ultrapure water (Optima™ LC/MS Grade, Fisher Chemical™).

### Protein extraction and peptide preparation

We prepared samples for metaproteomics analysis at two time-points: after storing the samples for 1 week and after 4 weeks. We removed the storage solutions from the samples by centrifugation at 21,000 × g for 5 min and then resuspended the samples in 400 μl of SDT lysis buffer [4% (w/v) SDS, 100 mM Tris-HCl pH 7.6, 0.1 M DTT]. Cells were lysed by bead-beating in lysing matrix E tubes (MP Biomedicals) with a Bead Ruptor Elite (Omni International) for 5 cycles of 45 sec at 6.45 m/s with 1 min dwell time between cycles, followed by heating at 95° C for 10 min. The lysates were centrifuged for 5 min at 21,000 × g to remove cell debris. We prepared peptides according to the filter-aided sample preparation (FASP) protocol described by [61]. All centrifugations mentioned below were performed at 14,000 × g. Samples were loaded onto 10 kDa MWCO 500 μl centrifugal filters (VWR International) by combining 60 μl of lysate with 400 μl of Urea solution (8 M urea in 0.1 M Tris/HCl pH 8.5) and centrifuging for 30 min. This step was repeated twice until the filter capacity was reached. Filters were washed twice by applying 200 μl of urea solution followed by 40 min of centrifugation. 100 μl IAA solution (0.05 M iodoacetamide in Urea solution) was then added to filters for a 20 min incubation followed by centrifugation for 20 min. The filters were washed three times with 100 μl of urea solution and 20 min centrifugations, followed by buffer exchange to ABC (50 mM Ammonium Bicarbonate). Buffer exchange was accomplished by adding 100 μl of ABC and centrifuging three times followed by centrifugation for 20 min. Tryptic digestion was performed by adding 0.85 μg of MS grade trypsin (Thermo Scientific Pierce, Rockford, IL, USA) in 40 μl of ABC to the filters which and incubating for 16 hours in a wet chamber at 37° C. The tryptic peptides were eluted by adding 50 μl of 0.5 M NaCl and centrifuging for 20 min. Peptide concentrations were determined with the Pierce Micro BCA assay (Thermo Fisher Scientific) following the manufacturer’s instructions.

### LC-MS/MS

Samples were analyzed by 1D-LC-MS/MS using a published method [62] with several modifications. The samples were blocked and randomized according to Oberg and Vitek’s method [63] to control for batch effects. For each sample, 600 ng of tryptic peptides were loaded with an UltiMate™ 3000 RSLCnano Liquid Chromatograph (Thermo Fisher Scientific) in loading solvent A (2 % acetonitrile, 0.05 % trifluoroacetic acid) onto a 5 mm, 300 μm ID C18 Acclaim® PepMap100 pre-column and desalted (Thermo Fisher Scientific). Peptides were then separated on a 75 cm × 75 μm analytical EASY-Spray column packed with PepMap RSLC C18, 2 μm material (Thermo Fisher Scientific) heated to 60 °C via the integrated column heater at a flow rate of 300 nl min^−1^ using a 140 min gradient going from 95 % buffer A (0.1 % formic acid) to 31 % buffer B (0.1 % formic acid, 80 % acetonitrile) in 102 min, then to 50 % B in 18 min, to 99 % B in 1 min and ending with 99 % B. Carryover was reduced by wash runs (injection of 20 μl acetonitrile with 99% eluent buffer B) between samples.

The analytical column was connected to a Q Exactive HF hybrid quadrupole-Orbitrap mass spectrometer (Thermo Fisher Scientific) via an Easy-Spray source. Eluting peptides were ionized via electrospray ionization (ESI). MS^1^ spectra were acquired by performing a full MS scan at a resolution of 60,000 on a 380 to 1600 m/z window. MS^2^ spectra were acquired using a data-dependent approach by selecting for fragmentation the 15 most abundant ions from the precursor MS^1^ spectra. A normalized collision energy of 25 was applied in the HCD cell to generate the peptide fragments for MS^2^ spectra. Other settings of the data-dependent acquisition included: a maximum injection time of 100 ms, a dynamic exclusion of 25 sec, and exclusion of ions of +1 charge state from fragmentation. About 60,000 MS/MS spectra were acquired per sample.

### Protein Identification Database

We constructed a protein sequence database for identifying proteins from the four main components of the sample: the host, wheat (the main component of mouse chow), the microbiota, and potential contaminants. Protein sequences of the mouse host, *Mus musculus*, were downloaded from Uniprot (https://www.uniprot.org/proteomes/UP000000589). Protein sequences of wheat, *Triticum aestivum,* were downloaded from Uniprot (https://www.uniprot.org/proteomes/UP000019116). For the microbiota sequences, a public database constructed by Xiao et al. [64] was used. While the use of such a reference database is not recommended for studies that address specific biological questions, as it has been shown that such reference databases can lead to lower identification rates and species and protein miss assignments [41, 65], it is sufficient for determining the overall effects of sample preservation and preparation methods. The database from Xiao et al. contains ~2.6 million “non-redundant” genes from metagenomic sequencing of fecal material from 184 mice. The corresponding annotated protein sequences were downloaded from http://gigadb.org/dataset/view/id/100114/token/mZlMYJIF04LshpgP. The taxonomy (available as a separate file) was integrated into the string of the sequence descriptions using the join command in Linux. Most (67.8%) of the sequences were assigned a taxonomy at the phylum level, and 9.8% of the sequences were assigned at the genus level [64]. Initial analyses suggested the presence of sequences that were too similar for adequate discrimination in the downstream workflow so the protein sequences were clustered with an identity threshold of 95% using the CD-HIT tool [66]. About 8% of the sequences were combined into clusters, while the remaining ~92% remained as individual sequences. Also included in the database were sequences of common laboratory contaminants (http://www.thegpm.org/crap/). The database contained a total of 2,396,591 protein sequences and is included with the PRIDE submission for this study (PXD024115).

### Protein identification and quantification

For peptide and protein identification, MS data were searched against the above-described database using the Sequest HT node in Proteome Discoverer version 2.3.0.523 (Thermo Fisher Scientific) with the following parameters: digestion with trypsin (Full), maximum of 2 missed cleavages, 10 ppm precursor mass tolerance, 0.1 Da fragment mass tolerance and maximum 3 equal dynamic modifications per peptide. We considered the following dynamic modifications: oxidation on M (+15.995 Da), carbamidomethyl on C (+57.021 Da), and acetyl on the protein N terminus (+42.011 Da). Peptide false discovery rate (FDR) was calculated using the Percolator node in Proteome Discoverer and only peptides identified at a 5% FDR were retained for protein identification. Proteins were inferred from peptide identifications using the Protein-FDR Validator node in Proteome Discoverer with a target FDR of 5%. From the generated multiconsensus dataset, we removed contaminant (cRAP sequences) and low confidence proteins (> 5 % FDR) and kept proteins that were identified by at least 1 protein unique peptide. To decrease the number of “one-hit wonder” proteins, we removed proteins that were not detected in a total of at least 3 samples (n = 47 samples in total), which is the minimum number of replicates in one treatment and time-point. The dataset contained 6,086 proteins after applying these filtering steps. The data were then normalized by calculating normalized spectral abundance factors (NSAFs, [67]) and multiplied by 100, to give the relative protein abundance in %.

### Quality assessment and outlier analysis

We assessed data quality by first inspecting raw data in the Xcalibur™ Software (Thermo Fisher Scientific), then by comparing the number of peptide spectrum matches (PSMs), peptides, proteins, and protein groups identified in each sample individually. We tested for statistical significance using a Student’s t-test (2-tailed, equal variability, FDR of 0.05). Samples in the dataset had on average 25,220 +/− 4.954 proteins identified at 5% FDR and 3,499 +/− 662 identified protein groups. Assuming the numbers of proteins identified per sample were normally distributed data, about 99.7% of the samples in the dataset were expected to have at least 10,358 detected proteins and 1,513 protein groups. These numbers correspond to the means stated above minus three standard deviations. One ethanol-preserved sample (“E6”) was deemed an outlier and was removed from the dataset because it had only 767 proteins and 84 protein groups. We suspect the protein extraction for that particular sample failed due to leaks in the filter unit during sample preparation.

### Data analyses

To investigate the degree of overlap in protein identifications between treatments, we used the filtered dataset of 6,086 proteins. If a protein was detected in at least one sample of a treatment within this dataset, it was counted as identified in that treatment. We imported the accession codes of the identified proteins into Venny 2.1 [68] to create Venn diagrams representing the overlap between treatments in terms of identified proteins.

To identify differentially abundant proteins between treatments that are statistically significant, we performed a centered-log-ratio (CLR) transformation in R (version 4.0.2, compositions_2.0-1 package)[69] on peptide spectrum matches (PSMs) before performing statistical tests. We added 1 to every PSM value before performing the CLR transformation to protect against issues with missing values. Although CLR-normalized counts lose interpretability, CLR is a method better suited for statistical analyses of compositional data such as metaproteomics data [70, 71]. Pairwise comparisons of all treatments were performed in the Perseus software platform (version 1.6.12.0) [42] using a Student’s T-test corrected for multiple hypothesis testing with a permutation-based FDR of 5% (S0=0.1, both sides, not paired).

We used a principal component analysis (PCA) to visualize how samples separate or cluster based on relative protein abundances. We performed the analysis in the Perseus software platform (version 1.6.12.0) [42] on the CLR-transformed dataset described above.

We investigated whether the preservation treatment affected the measured abundances of specific taxa by comparing the relative biomass contributions of the taxa. Biomass contributions of specific taxa were assessed at the phylum and genus levels using the method described in Kleiner et al. [32]. Briefly, proteins were filtered for at least 2 protein unique peptides to increase the confidence in taxonomic identifications, and PSMs summed by taxon were used to estimate the biomass contribution of each taxon in the metaproteomes.

We investigated whether preservation treatments were biased towards proteins with specific biochemical characteristics such as isoelectric point (pI), molecular weight, or presence of transmembrane domains. We retrieved the pI and molecular weight associated with each identified protein from Proteome Discoverer and detected transmembrane domains by searching sequences of the identified proteins on the TMHMM 2.0 Server [72]. Then, we compared the distributions of these properties in each treatment as histograms with defined ranges.

Lastly, we assessed the amount of within-treatment variation using linear regression scatterplots in R (version 4. 0. 2; psych_2.1.3 package). We fit the scatterplots onto the percent normalized spectral abundance factors (%NSAFs) for each pair of replicates that received the same treatment (n=4, except for the ethanol treatment time point 4 weeks: n=3 because sample E6 was removed). We then compared the Pearson correlation coefficients.

### Data availability

The mass spectrometry metaproteomics data and protein sequence database were deposited to the ProteomeXchange Consortium via the PRIDE [73] partner repository with the dataset identifier PXD024115 [Reviewer Access at https://www.ebi.ac.uk/pride/login User: Password: WVJJYH44]

## Results

A fecal master mix (homogenate) was prepared from fecal samples of healthy adult conventional C57BL/6 mice. Aliquots of the master mix were randomized and then preserved using different methods. After 1 week and 4 weeks of storage, proteins were extracted and analyzed by LC-MS/MS. Preservation methods were assessed based on the amount of variability between replicates, and the degree of bias when compared to the other methods, particularly flash-freezing.

### Minimal differences in total numbers of identified features for co-extracted samples

We compared the number of peptide spectrum matches (PSMs), peptides, proteins, and protein groups identified at a false discovery rate (FDR) of 5% between the different treatments and time-points (**Figure 1**) to determine whether the preservation treatment impacted the number of features identified. Samples that were preserved for 1 week and co-extracted as part of the first extraction batch did not significantly differ in their total counts, regardless of the preservation method. The numbers of features for the 4-week time-point (2nd extraction batch) and 1-week time-point differed slightly but differences did not test significant except for the samples preserved at −80° C in RNA*later*™ (“RF” samples). This difference is likely due to batch effects in sample preparation and peptide quantification via microBCA assay because the 1-week and 4-week samples were prepared separately. At the 4-week time-point, flash-frozen (FF) samples preserved for 4 weeks at −80° C were significantly lower in their total counts compared to RF samples or samples preserved in the NAP buffer (N) or autoclaved NAP buffer (AN) at room temperature.

**Figure 1.**
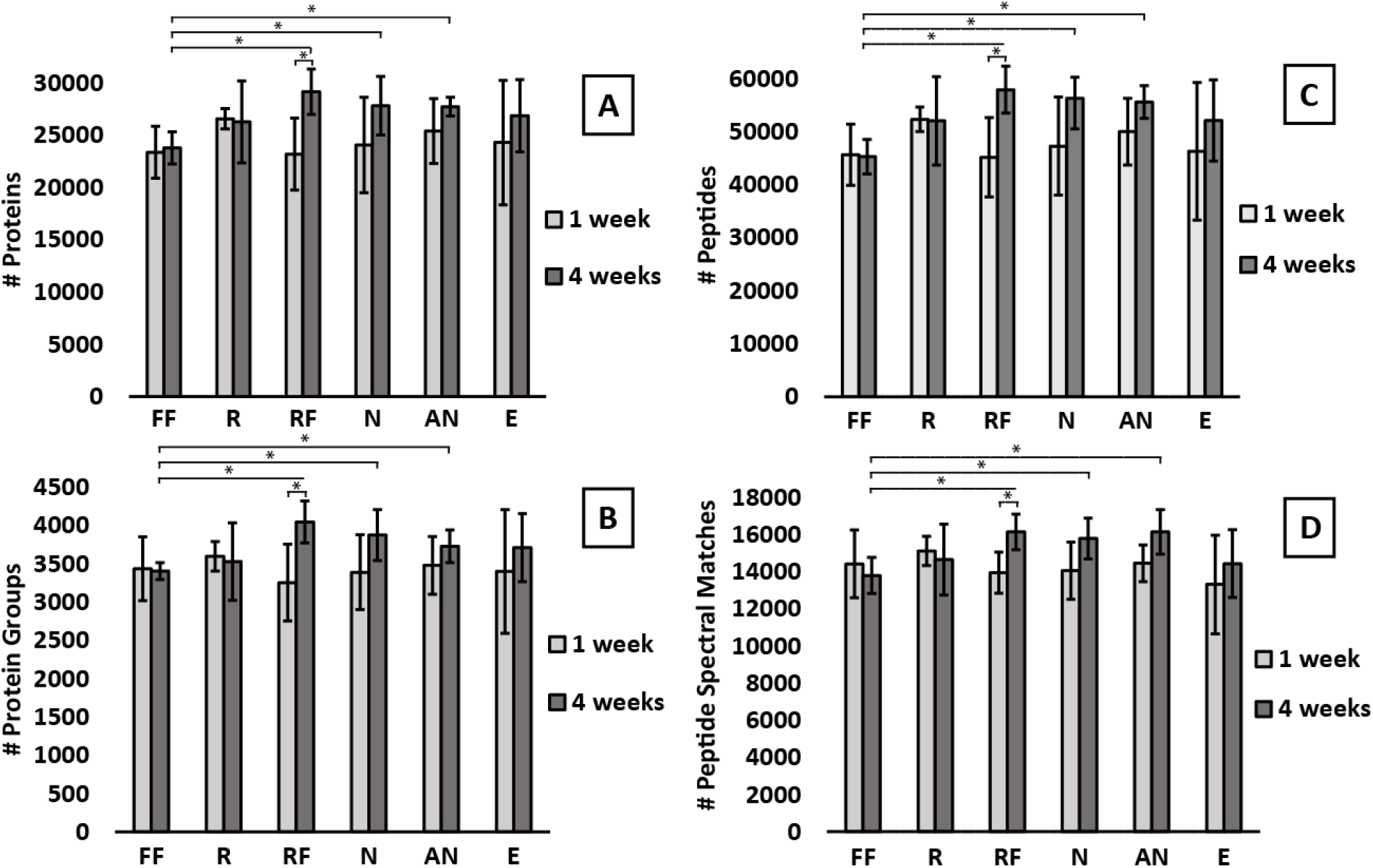
There were no significant differences in total numbers of PSMs, peptides, proteins, and protein groups between samples co-extracted after 1 week of preservation and only minimal differences existed in samples co-extracted after 4 weeks. FF = Flash-freezing, R = RNAlater, RF = RNAlater + flash freezing, N = NAP buffer, AN = Autoclaved NAP buffer, E = 95% Ethanol. 1 week = preserved for one week and first extraction batch. 4 weeks = preserved for 4 weeks and second extraction batch. Bars represent the arithmetic mean (n = 4 for all except E - 4 weeks where n = 3). Error bars represent standard deviation. Asterisks indicate statistical significance (t-test, p-value < 0.05). **A)** Total proteins identified at 5% FDR include the microbial, host, and dietary proteins. **B)** Total protein groups identified at 5% FDR. **C)** Total peptides identified at 5% FDR. **D)** Total peptide spectrum matches (PSMs) identified at 5% FDR.

### Treatments shared over 76% of protein identifications

Comparing and quantifying the proteins identified by multiple treatments showed that most proteins were detected in every treatment. **Figure 2A** shows a Venn diagram of the four most distinct treatments in terms of physical and chemical properties: R, E, FF, and N. Each of these 4 treatments produced metaproteomes that identified the same 4,641 proteins (76.3% of the dataset) and uniquely identified ~1% of proteins. In the E treatment, 281 proteins or 4.6% of the protein identifications were not detected; this was the largest proportion of undetected proteins, followed by the FF treatment that did not detect 198 proteins or 3.3% of the protein identifications. **Figure 2B** represents the overlap of proteins between chemically similar treatments: R, RF, N, and AN. Each of these 4 treatments produced metaproteomes that identified the same 4,878 proteins (80.4% of the dataset) and uniquely identified ~1% of proteins. The RF treatment was the most distinct of the comparison shown in **Figure 2B**(R, RF, N, AN) with 74 unique protein identifications (~1.2%) that were not detected in the other treatments.

**Figure 2.**
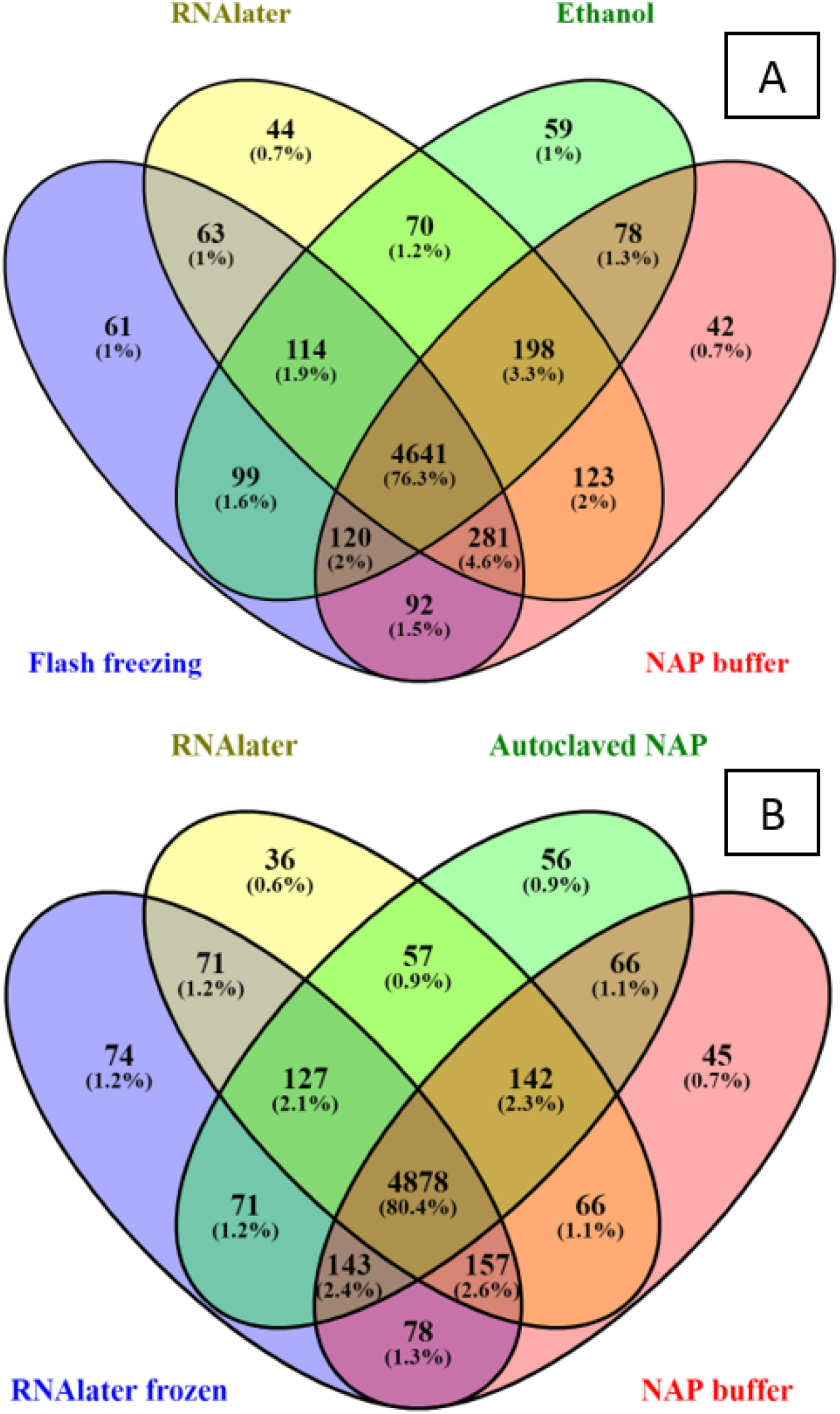
Over 76% of microbial, host and dietary protein identifications overlapped between treatments. Replicates of both time points were combined (n = 8 samples per treatment, except for Ethanol where n = 7). Proteins were included if they were identified with FDR <5%, at least 1 protein unique peptide, and present in at least 3 samples in the whole dataset. **A)** Comparison of treatments that differed most in terms of physical/chemical properties: Flash freezing (FF), RNAlater (R), 95% Ethanol (E), and NAP buffer (N). **B)** Comparison of the chemically similar treatments (RNAlater™ and RNAlater-like treatments): RNAlater (R), RNAlater frozen (RF), NAP buffer (N), Autoclaved NAP (AN).

### Relative protein abundances were highly similar between all treatments except for the ethanol treatment

Principal component analysis (PCA), performed on the Centered-Log-ratio (CLR) transformed dataset of relative protein abundances, showed that the ethanol-preserved samples clustered together and clearly separated from samples of all other treatments (**Figure 3A**). If the preservation treatment did not affect protein profiles, we would expect to see no clustering, but rather the samples would be randomly distributed over the PCA plot.

**Figure 3.**
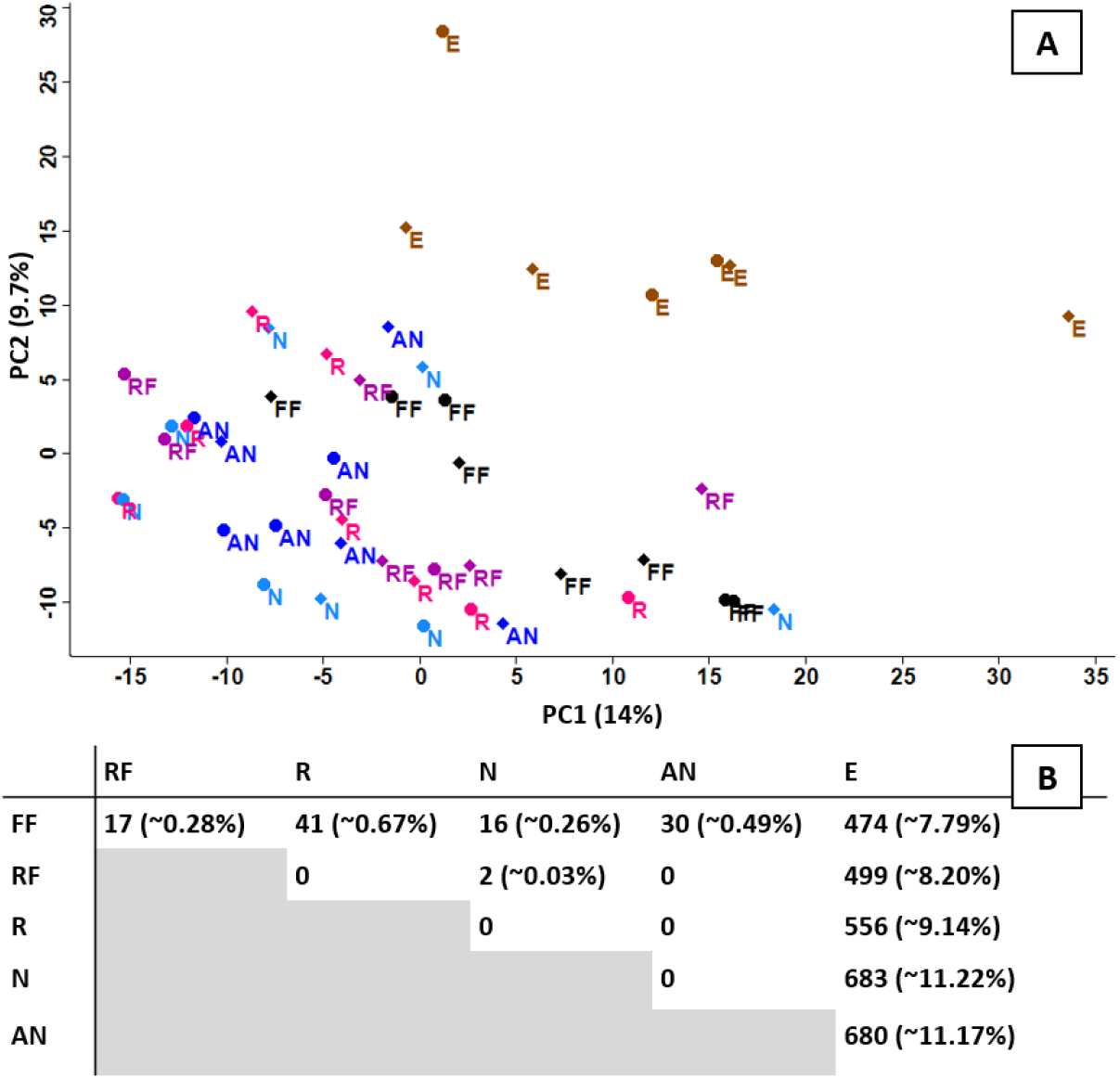
Ethanol preserved samples were distinct from all other samples in their protein abundance profiles. **A)** Principal component analysis (PCA) of the relative protein abundances from each sample (CLR-transformed). E = ethanol; FF = flash-freezing; R = RNA*later*™; RF = RNA*later*™ and flash-freezing; N = NAP buffer; AN = autoclaved NAP buffer. Diamonds = 1-week; Circles = 4-weeks. **B)** Number of significant differences between each treatment (two-sided t-test, FDR of 0.05 and S0 of 0.1). A significant difference represents one protein that is more abundant in one treatment over the other for each paired comparison (refer to Additional file 2 for directionality). Percentages in parentheses indicate the percentage of significant proteins out of the total proteins considered (n = 6,086).

We then compared CLR-transformed relative protein abundances between treatments using t-tests corrected for multiple hypothesis testing (two-sided, FDR of 0.05 and S0 of 0.1) to identify proteins that significantly differed in abundance based on the preservation treatment (Figure 3B). We found no significant differences between RNA*later*™(R) and the NAP buffer (N) or between the NAP buffer (N) and the autoclaved NAP buffer (AN). The proteins not shared between these treatments (Fig 2A) were sparse and lowly abundant proteins that were not significantly different from an undetected protein **(Additional file 2)**. The ethanol treatment was the most distinct treatment with, on average, about 9.5% of the proteins significantly differing in abundance, with 247 proteins being less abundant, as compared to the other treatments **(Additional file 2).** Flash-frozen samples and the samples preserved in RNAlater or RNAlater-like solutions produced metaproteomes that differed only minimally (< 1% of proteins with different abundances).

### Within-treatment variability of relative protein abundances was low

We assessed the amount of within-treatment variability in terms of quantified protein abundances by fitting linear scatter plots for all replicates against all replicates and evaluating the Pearson correlation coefficients (**Additional file 6: Figures S1 - 6**). The means of the Pearson correlation coefficients (**Table 1**) showed high correlation between replicates, indicating within treatment variability was low for all treatments.

**Table 1.** Linear correlation of replicates. Mean Pearson coefficient of the linear correlation between replicates of the same treatment and preservation duration (n = 4, except for E 4 weeks: n = 3). We fit a linear model for each pair of samples within a treatment and time point in R (version 4. 0. 2; psych_2.1.3 package) using the dataset of percent normalized spectral abundance factors (%NSAFs).

|  |  |  |
| --- | --- | --- |
|  | 1 week | 4 weeks |
| **NAP buffer** | 0.937 +/- 0.024 | 0.963 +/- 0.006 |
| **Autoclaved NAP buffer** | 0.965 +/- 0.008 | 0.958 +/- 0.02 |
| **RNAlater™** | 0.957 +/- 0.01 | 0.957 +/- 0.017 |
| **RNAlater™ + Flash-freezing** | 0.938 +/- 0.034 | 0.967 +/- 0.008 |
| **Flash Freezing** | 0.961 +/- 0.009 | 0.943 +/- 0.019 |
| **95% Ethanol** | 0.957 +/- 0.01 | 0.95 +/- 0.015 |

### Small but significant differences in the taxonomic composition of the metaproteomes based on the preservation method

The relative taxonomic composition of the samples in terms of proteinaceous biomass contribution was consistent across replicates and preservation treatments (**Additional text and additional. fig. S7**). The biomass contribution is shown per phylum in **Fig. 4A** and per genus in **Fig. 4B** for the most abundant genera: *Clostridium, Eubacterium, Butyrivibrio, Lactobacillus, Turicibacter, Blautia, Roseburia,* and *Coprococcus*. The abundances of specific taxa significantly differed at the phylum and genus levels. At the phylum level, Firmicutes was overrepresented in the ethanol-preserved samples as compared to the flash-frozen and NAP-buffer preserved samples. At the genus level, *Clostridium* and *Blautia* were subtly but significantly overrepresented in the ethanol-preserved samples as compared to all other treatments (t-test, paired, 2-tailed, p < 0.05). Furthermore, NAP buffer and Ethanol-preserved samples differed in their representation of the genus *Butyrivibrio*, and RNAlater and Ethanol-preserved samples differed in their representation of the genus *Roseburia*.

**Figure 4.**
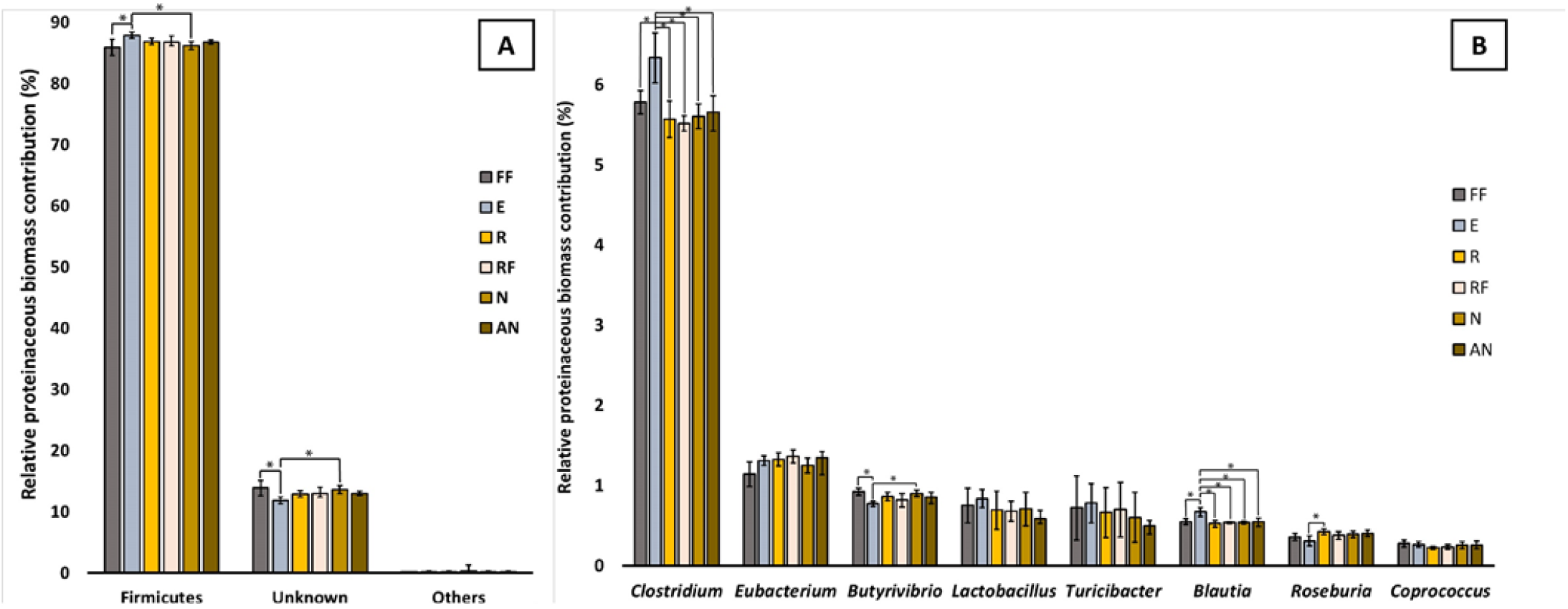
Small but significant differences in the representation of microbial taxa in the metaproteomes based on preservation method. Bars represent the mean percent proteinaceous biomass for each taxon at the phylum level (A) or the genus level (B). Error bars represent the standard deviation (n=8, except n=7 for the Ethanol treatment). Asterisks represent statistical significance (t-test, paired, 2-tailed, p < 0.05). E = ethanol; FF = flash-freezing; R = RNAlater; RF = RNAlater and flash-freezing; N = NAP buffer; AN = autoclaved NAP buffer. The eight most abundant genera are displayed in the figure. Percentages are low because genus-level taxonomy could be assigned for 11.1 +/− 0.53 % (n = 47) of the total proteinaceous biomass in our samples, distributed over 28 different microbial genera.

### The preservation methods did not bias towards specific biochemical properties of proteins

We investigated whether the preservation treatment biased towards or against proteins with a specific isoelectric point (pI), molecular weight **(Additional Data 3)**, or transmembrane domains **(Additional Data 4)** by comparing the distributions of these properties in each treatment. Distributions did not differ between treatments, indicating no bias **(Fig. 5)**.

**Figure 5.**
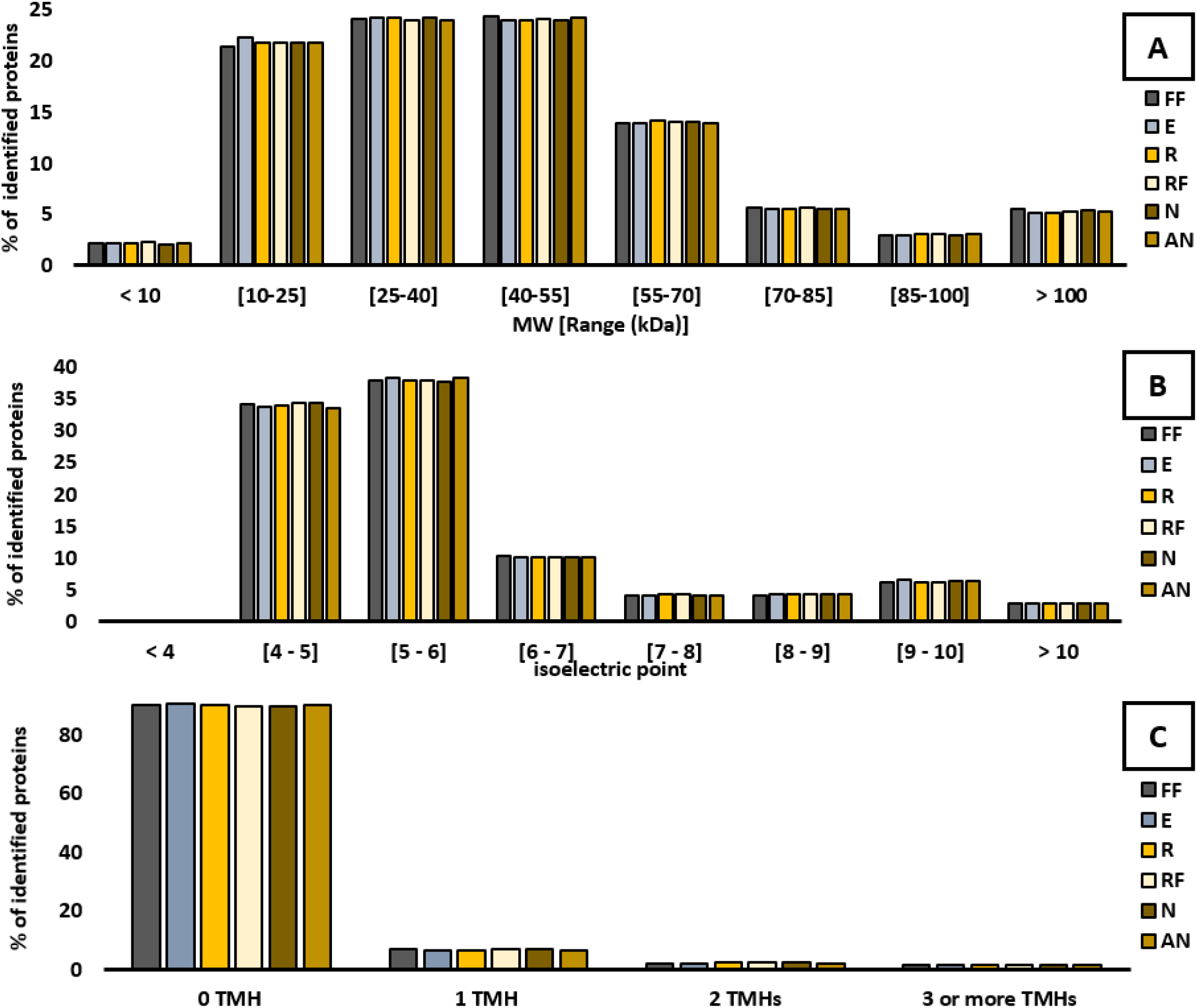
Distribution of biochemical properties of identified proteins. A) molecular weight (kDa) B) isoelectric point C) number of predicted transmembrane helices. Bars represent the proportion (%) of identified proteins belonging in each range. E = ethanol; FF = flash-freezing; R = RNAlater; RF = RNAlater and flash-freezing; N = NAP buffer; AN = autoclaved NAP buffer.

## Discussion

This study evaluated the effects of different preservation treatments on the metaproteomes of intestinal microbiome samples to identify a preservation method suitable to use when flash-freezing is not an option. The data show that the metaproteomes of samples preserved at room temperature while immersed in RNA*later*™ or RNAlater-like solutions (NAP buffer and autoclaved NAP buffer) were highly similar to the metaproteomes of samples preserved by flash-freezing and storage at −80°C, with only negligible differences. On the other hand, samples preserved by immersion in 95% ethanol differed substantially from the flash-frozen samples and other samples. Because methods sharing the largest number of discoveries/values with most of the other methods tested may be more likely to produce valid results [74], our results suggest that the 95% ethanol treatment creates the largest bias in the metaproteomes. In contrast, the flash freezing, RNAla*ter*™, and RNAlater-like treatments are most likely to represent the protein profiles at the time of collection accurately. The majority of the differences of the ethanol treatment were found at the protein abundance level. Roughly 9.5% of all the proteins in the ethanol preserved samples differed in abundance from all other samples, suggesting that ethanol preservation could strongly influence study results. Taxonomic abundances also differed under the ethanol treatment with the genera *Clostridium* and *Blautia* being overrepresented compared to all other treatment samples. However, although ethanol preservation introduces biases, within-treatment variability was low in ethanol-preserved samples. These results suggest that ethanol may be appropriate in some studies if it is used consistently.

While we tested a diversity of popular preservation methods, there are potentially other storage solutions and methods that could be used in addition to or instead of those tested and described herein. There are, for example, a range of commercially available “microbiome” storage solutions designed for preservation of fecal material for amplicon sequencing. These solutions could potentially also be used for metaproteomics, however, their compatibility with the proteomic workflow and quality of preservation would have to be carefully tested, particularly as compatibility issues could arise if a preservation reagent is incompatible with standard proteomic workflows. For example, some preservatives contain guanidinium chloride, which forms a solid if it contacts SDT lysis buffer [4% (w/v) SDS, 100 mM Tris-HCl pH 7.6, 0.1 M DTT] that is used in many metaproteomics workflows.

Our results concur with the study by Saito et al. [55], which investigated sample preservation effects on the proteome of the cyanobacterium *Synechococcus* WH8102. They found RNA*later*™ to be effective at preserving the proteome of a pure culture when compared to frozen storage. Here, we demonstrated that RNA*later*™ can also be effective at preserving the metaproteomes of complex microbiome samples while minimizing storage effects. Furthermore, it appears that using the cost-effective RNAlater-like solution “NAP buffer” is a suitable alternative to the commercial RNA*later*™ solution. Menke et al. [58] previously demonstrated that the NAP buffer effectively preserves the DNA of intestinal microbiome samples. Here we demonstrated that the NAP buffer effectively preserved proteins of intestinal microbiome samples and did not affect protein abundances when compared to samples preserved in RNA*later*™. Furthermore, autoclaving the NAP buffer did not significantly affect the metaproteomes, suggesting that an autoclaved NAP buffer could be used in studies that require sterile material (e.g., gnotobiotic isolators).

## Conclusions

This study evaluated the effects of different preservation treatments on the metaproteomes of intestinal microbiome samples. Based on our results, we recommend preserving intestinal microbiome samples by freezing, or in RNAlater™, or an RNAlater-like solution before metaproteomics analyses. The consistent use of these methods appears to minimize storage effects and thus improve the reliability of metaproteomics studies of the intestinal microbiome.

## Declarations

### Ethics Approval

The protocols for husbandry and experimentation of all mice used in this study were approved by the Institutional Animal Care and Use Committee at North Carolina State University (Institution reference: D16-00214).

### Consent for publication

Not applicable

### Competing interests

The authors declare no competing interests.

### Funding

This work was supported by the National Institute Of General Medical Sciences of the National Institutes of Health under Award Number R35GM138362 and the Foundation for Food and Agriculture Research (FFAR) Grant ID: 593607

### Authors’ contributions

A.M. and M.K. designed the study. A.M. performed the experiments. A.M. and M.K. analyzed the data and wrote the manuscript.

## Acknowledgments

We thank Deniz Durmusoglu for providing the fresh mouse fecal samples, Dr. Fernanda Salvato for providing A.M. with extensive metaproteomics training, Ibrahim Al’Abri and Alexandria Bartlett for providing mouse fecal samples for prior optimization rounds of the study, Marlene Jensen, Clara Tang, and Amelia Ellis who provided assistance in the lab, everyone in the Kleiner Lab for discussing the experimental design and results, and Dr. Heather Maughan for feedback on the manuscript. We made all LC-MS/MS measurements in the Molecular Education, Technology, and Research Innovation Center (METRIC) at NC State University.

